# Persistence of cultivar alleles in wild carrot (*Daucus carota* L.) populations in the United States

**DOI:** 10.1101/2022.12.05.519198

**Authors:** Fernando Hernández, Luciano Palmieri, Johanne Brunet

## Abstract

Cultivated species and their wild relatives often hybridize in the wild and crop-wild hybrids can survive and reproduce in some environments. However, it is unclear whether crop alleles are permanently incorporated into the wild genomes in the long run or whether they are purged by natural selection. This question is key to accurately assessing the risk of escape and spread of cultivar genes into wild populations. Here, we use genomic data and population genomic methods to study hybridization and introgression between cultivated and wild carrots (*Daucus carota* L.) in the United States. We used single nucleotide polymorphisms (SNPs) obtained via genotyping by sequencing for 450 wild individuals from 29 wild georeferenced populations in seven states and 144 cultivars from the United States, Europe, and Asia. Cultivated and wild carrots formed two well differentiated groups, and evidence of crop-wild admixture was detected in several but not all the wild populations in the United States. Two regions were identified where cultivar alleles were introgressed into wild carrots: California and the Nantucket Island, in Massachusetts. In these areas, we found no support for adaptive (or maladaptive) introgression, instead, most crop alleles seemed to be neutral. Surprisingly, there was no evidence of introgression in some populations with a long-known history of sympatry with the crop, suggesting that post-hybridization barriers might prevent introgression in some areas. Further studies are needed to better delineate the geographic patterns of introgression, but our results support the introgression and persistence of cultivar genes in wild carrot populations.

## INTRODUCTION

Gene flow between crop species and their wild relatives has played an important role in the evolution of both wild and cultivated plants (Ellstrand & Rieseberg, 2016). Introgression of wild alleles into cultivated germplasm constitutes an important source of new genetic variation for crop breeding (Hajjar & Hodgkin, 2007; Prohens et al., 2017; Renzi et al., 2022). However, introgression from crops to nearby wild populations can have a negative impact on wild populations by decreasing genetic diversity and even causing local extinction of the recipient populations (Hegde, Nason, Clegg, & Ellstrand, 2006; Todesco et al., 2016); it may also facilitate the evolution of agricultural weeds (Ellstrand et al., 2010; Le Corre, Siol, Vigouroux, Tenaillon, & Délye, 2020; Schierenbeck & Ellstrand, 2009).

When crop and wild plants grow in sympatry and have overlapping flowering periods, hybridization may occur in both directions. Even if the crop and wild hybridize, the probability of successful hybridization leading to permanent gene introgression depends on the relative fitness of the hybrid to non-hybrid progeny (Mercer, Andow, Wyse, & Shaw, 2007; Snow, Moran-Palma, Rieseberg, Wszelaki, & Seiler, 1998). The reduced fitness of F1 hybrids is a major ecological barrier for the introgression of crop alleles into wild populations. However, the strength of this barrier varies with the environmental and ecological contexts (Mercer et al., 2007; Presotto, Hernández, & Mercer, 2019) and it is expected to weaken as the hybrids backcross with wild plants (Gutierrez, Cantamutto, & Poverene, 2011; Hooftman, De Jong, Oostermeijer, & Den Nijs, 2007; Presotto, Ureta, Cantamutto, & Poverene, 2012). Moreover, specific crop alleles that increase fitness in agricultural environments (e.g., herbicide resistant alleles) can quickly introgress into wild populations, contributing to the evolution of aggressive weeds (Le Corre et al., 2020; Pandolfo et al., 2018; Singh et al., 2017).

The recent advances in genome-editing techniques, such as CRISPR/Cas9, and their use in crop improvement have rekindled the interest in crop-wild gene flow (Chen, Wang, Zhang, Zhang, & Gao, 2019; Puchta, 2017). The number of crops being genetically modified (GM), and the number of GM cultivars of a crop being generated is expected to rise sharply in the near future as a result of these technologies, and cultivated carrot, *Daucus carota* L. subsp. *sativus*, is an example. Klimek-Chodacka *et al*. (2018) reported the first successful site-directed mutagenesis with CRISPR/Cas9 in carrot, paving the way for the use of genome-editing tools in carrot breeding. Since wild and cultivated carrots belong to the same species, share pollinators, and are highly outcrossed, hybridization is very likely in sympatry (Rong, Janson, Umehara, Ono, & Vrieling, 2010). Moreover, cultivated carrot is grown worldwide and wild carrot is present in temperate areas throughout the world. Wild carrot has been declared invasive in many states in the USA (Mandel and Brunet 2019). Thus, there is serious concern about the evolution of agricultural carrot weeds and increased invasiveness of wild carrot due to crop-wild hybridization from genome-edited carrot cultivars. This concern is reinforced by the detection of natural hybrids in regions where wild carrot populations and crop fields occur in close proximity (Mandel, Ramsey, Iorizzo, & Simon, 2016; Palmieri, Ellison, Senalik, Simon, & Brunet, 2019). Moreover, in some environments crop-wild hybrids can survive and reproduce almost as well as wild individuals (Hauser & Shim, 2007; Magnussen & Hauser, 2007; Rong et al., 2010). However, even if hybridization between cultivated and wild carrots has been detected, it is unclear whether hybridization leads to permanent gene introgression (Mandel et al., 2016; Palmieri et al., 2019). Recent studies in crop wild hybrid zones for different species, including sunflower (Mondon, Owens, Poverene, Cantamutto, & Rieseberg, 2018), and Lima bean (Heredia-Pech et al., 2022), failed to detect permanent introgression of crop alleles into wild populations, suggesting that even when hybridization takes place and hybrids survive, crop alleles may be purged out by natural selection in the long run. The potentially negative consequences of introgression of genome-edited genes in wild carrots warrants an assessment of the risk of escape and spread of cultivar genes into wild carrot populations.

Phenotypic detection of hybrids can be difficult. In species where hybrids can be identified morphologically, off-type plants can be used to infer hybridization and introgression in the field, e.g., in sunflower (Ureta, Cantamutto, Carrera, Delucchi, & Poverene, 2008) and oilseed rape (Ureta et al., 2017). However, phenotypic differences between hybrid and non-hybrid plants tend to disappear after a few backcrosses with wild populations, and in some species, hybrids cannot be easily identified morphologically (Zalapa, Brunet, & Guries, 2010). Phenotypic differences between wild and cultivated carrot reside mostly on root traits, i.e., cultivated carrots have been selected for colored (orange, yellow, or purple), sweet and palatable roots (Ellison et al., 2018; Iorizzo et al., 2016). No obvious differences exist between crop and wild in aboveground traits, which challenges the visual identification of crop-wild hybrids. However, molecular markers from genome sequencing can provide a useful tool for identifying crop-wild hybrids in recent and advanced generations of backcrossing (Muller, Latreille, & Tollon, 2011; Palmieri et al., 2019; Torres Carbonell, Ureta, Pandolfo, & Presotto, 2020).

Examining introgression can be challenging even with the use of genomic data. The main focus of most crop-wild gene flow studies is to assess the risk of allele escape from current cultivars to nearby wild or weedy populations (Mondon et al., 2018; Palmieri et al., 2019; Pandolfo et al., 2016; Snow et al., 2010; Stewart, Halfhill, & Warwick, 2003). However, due to the relatively recent evolution of crops (Purugganan & Fuller, 2009), and the recurrent use of wild relatives in plant breeding (Hajjar & Hodgkin, 2007; Prohens et al., 2017), differentiating recent crop to wild gene flow from historical gene flow (e.g., during domestication), and from recent wild to crop introgression in plant breeding can be challenging with traditional approaches (Hibbins & Hahn, 2022; Martin & Amos, 2020; Payseur & Rieseberg, 2016). Recently, robust population genomic and phylogenomic methods have been developed to detect past hybridization events in genomes of contemporary samples, and to infer not only the presence of hybrids but also the amount, timing, and direction of gene flow (Bourgeois & Warren, 2021; Hibbins & Hahn, 2022; Martin & Amos, 2021; Payseur & Rieseberg, 2016). Though designed for speciation studies, these methods can be applied to crop-wild gene flow studies.

Here, we used genomic data and population genomic methods to study hybridization and introgression between cultivated and wild carrots in the United States. To reach this goal we (1) tested whether wild carrot populations show evidence of admixture with cultivated carrot; (2) evaluated geographic patterns of hybridization; and (3) studied the timing and direction of gene flow using state-of-the-art population genomic methods. Recent crop to wild introgression was detected in some but not all populations in the United States.

## MATERIALS AND METHODS

### Plant species

Cultivated and wild carrot belong to the same species, *Daucus carota* L. Cultivated carrot (*Daucus carota* L. subsp. *sativus*) is a common vegetable, grown worldwide mostly for root consumption (Iorizzo et al., 2013). Nikolai Vavilov (1992) proposed central Asia as the center of origin of cultivated carrots, where the species has been used as a root vegetable for at least 1000 years. Genetic studies have provided strong support for a single origin of cultivated carrot in central Asia, followed by dispersal of domesticated carrot across Asia and the Mediterranean region, and then to the Americas (Iorizzo et al., 2016, 2013; Rong et al., 2014). In the United States, cultivated carrot seed production occurs in areas where wild carrot is rare or absent to minimize contamination and maintain cultivar purity, however, carrots for root production often grow in sympatry with wild carrot populations (Mandel et al., 2016).

Wild carrot, or Queen Anne’s lace (*Daucus carota* L. subsp. *carota*) is the ancestor of cultivated carrot, and it is widely distributed across temperate regions of the world (Iorizzo et al., 2013). In the United States, wild carrot was introduced by European settlers, probably in the 17^th^ century (Iorizzo et al., 2013). It is distributed across the country and has been declared invasive in many states (CABI, 2022). Wild populations are commonly found in ruderal environments, such as roadsides, waste places, and abandoned crop fields (CABI, 2022).

### Sample collection

We collected samples from 450 individuals of Queen Anne’s lace from 29 georeferenced wild populations in seven states between 2007 and 2019 (Table 1); leaves were collected *in situ* for DNA extraction. For each site, three climatic variables (Table 1) were obtained from the BIOCLIM dataset using the *raster* R package (Hijmans, Cameron, Parra, Jones, & Jarvis, 2005; Hijmans & van Etten, 2012).

**Table 1.** Information of the wild populations used in this study. Sample size after filtering. BIO1: mean annual temperature (°C); BIO6: minimum temperature of the coldest month (°C); BIO12: annual precipitation (mm). Year is the year that samples were collected. Population size (Pop size) is the estimated number of individuals in the population at the time of collection.

| Population | Region | Sample size | Latitude | Longitude | BIO1 | BIO6 | BIO12 | Year | Pop size | Proximity to crop |
| --- | --- | --- | --- | --- | --- | --- | --- | --- | --- | --- |
| CA1 | California | 17 | 38.56 | -121.38 | 16.1 | 3.4 | 453 | 2011 | 100 | NO |
| CA2 | California | 20 | 38.89 | -121.01 | 15 | 1.7 | 947 | 2011 | 100 | NO |
| CA3 | California | 11 | 38.39 | -122.35 | 15 | 2.4 | 714 | 2019 | 1,000 | NO |
| CA4 | California | 11 | 37.47 | -122.43 | 12.9 | 5.5 | 630 | 2019 | 300 | NO |
| CA5 | California | 10 | 39.26 | -121.04 | 12.1 | -1.1 | 1433 | 2019 | 100 | NO |
| CO | Colorado | 11 | 39.70 | -104.87 | 9.9 | -8.8 | 415 | 2019 | 30 | NO |
| IA1 | Iowa | 10 | 42.45 | -92.10 | 8 | -14.3 | 874 | 2009 | 1,000 | NO |
| IA2 | Iowa | 13 | 42.00 | -93.61 | 8.9 | -13.6 | 835 | 2009 | 100 | NO |
| IA3 | Iowa | 10 | 42.33 | -90.87 | 8.1 | -13.7 | 891 | 2019 | 10,000 | NO |
| IA4 | Iowa | 11 | 41.98 | -93.25 | 8.3 | -13.8 | 860 | 2019 | 10,000 | NO |
| IA5 | Iowa | 10 | 41.60 | -93.79 | 9.4 | -12.5 | 837 | 2019 | 10,000 | NO |
| MA1 | Nantucket | 10 | 41.26 | -70.18 | 9.8 | -4.5 | 1101 | 2015 | 120 | YES |
| MA2 | Nantucket | 10 | 41.27 | -70.10 | 9.7 | -4.3 | 1096 | 2015 | 50 | YES |
| MA3 | Nantucket | 9 | 41.30 | -70.04 | 9.7 | -4.4 | 1099 | 2015 | 50 | YES |
| MA4 | Nantucket | 10 | 41.29 | -70.13 | 9.7 | -4.5 | 1100 | 2015 | 70 | YES |
| MN | Minnesota | 10 | 43.69 | -96.21 | 7.1 | -16.3 | 677 | 2019 | 250 | NO |
| SD | South Dakota | 10 | 44.45 | -103.86 | 7.6 | -11.3 | 545 | 2019 | 250 | NO |
| WI1 | Wisconsin | 10 | 43.06 | -89.55 | 7.4 | -14.1 | 826 | 2017 | 100 | YES |
| WI2 | Wisconsin | 18 | 43.06 | -89.54 | 7.4 | -13.9 | 823 | 2017 | 250 | YES |
| WI3 | Wisconsin | 20 | 43.06 | -89.54 | 7.4 | -13.9 | 823 | 2017 | 1,000 | YES |
| WI4 | Wisconsin | 19 | 43.07 | -89.54 | 7.4 | -13.9 | 822 | 2017 | 750 | YES |
| WI5 | Wisconsin | 19 | 43.13 | -89.44 | 7.8 | -13.6 | 797 | 2017 | 1,000 | NO |
| WI6 | Wisconsin | 20 | 43.08 | -89.80 | 7.7 | -14.1 | 817 | 2017 | 1,000 | NO |
| WI7 | Wisconsin | 14 | 43.03 | -89.63 | 7.6 | -14.0 | 824 | 2017 | 750 | NO |
| WI8 | Wisconsin | 12 | 43.10 | -89.52 | 7.6 | -13.8 | 805 | 2017 | 750 | NO |
| WI9 | Wisconsin | 17 | 43.08 | -89.43 | 7.8 | -13.5 | 801 | 2007 | 1,000 | NO |
| WI10 | Wisconsin | 8 | 43.03 | -89.55 | 7.5 | -13.9 | 828 | 2010 | 1,000 | NO |
| WI11 | Wisconsin | 18 | 43.00 | -89.50 | 7.6 | -13.8 | 819 | 2007 | 5,000 | NO |
| WI12 | Wisconsin | 4 | 43.05 | -89.55 | 7.4 | -13.9 | 824 | 2007 | 5,000 | NO |

For cultivated samples, we included 144 cultivated accessions (hereafter cultivars) comprising inbred lines, commercial varieties, and landraces (Table S1). These 144 cultivars were classified by origin in CROP_US (15 cultivars grown in the United States), CROP_EAST (61 cultivars from middle East and Asia, including one sample from Ethiopia and two from Egypt), CROP_EUROPE (40 cultivars from western Europe), and CROP_OTHER (25 pre-breeding lines from the Wisconsin Breeding program, two cultivars from South Africa, and one from New Zealand). Cultivated accessions from the East, Europe, and other countries are deposited in the USDA germplasm bank (PI available in Table S1) and the pre-breeding lines from the Wisconsin Breeding Program were generously provided by Dr. Philipp Simon (United States Department of Agriculture-Agricultural Research Service).

### DNA extraction and sequencing

Total genomic DNA was extracted from lyophilized powdered leaf tissue using either a Qiagen DNeasy Mericon kit as in Palmieri et al. (2019) (samples extracted at the University of Wisconsin Biotechnology Center) or a Qiagen DNeasy Plant kit (samples extracted in the Brunet laboratory). Extractions performed with a DNeasy Plant kit followed the manufacture’s protocol with the following adjustments: samples were incubated overnight with proteinase K, and two successive elution steps were performed at the end with heated ddH2O (56°C) and 10 min incubation before centrifugation. The final DNA solution was treated with 5 μl RNAse A.

The majority of the 412 samples of Queen Anne’s lace (5 – 20 per population) and the pre-breeding lines from the Wisconsin breeding program were genotyped for this study. The other sequences had previously been generated (see Palmieri et al., 2019). Genotyping-by-sequencing (GBS) was performed as described by Elshire et al., (2011). DNA samples were digested with ApeKI, barcoded, and pooled for sequencing using 150 bp paired end reads on an Illumina NovaSeq 6000 platform. Library preparation and sequencing were performed at the University of Wisconsin-Madison Biotechnology Center DNA Sequencing Facility, in Madison, Wisconsin, USA.

### SNP calling and filtering

The resulting raw fastq files were demultiplexed, quality filtered, and assembled using the bioinformatics pipeline TASSEL-GBS v5.2.39 (Bradbury et al., 2007; Glaubitz et al., 2014). All sequences were aligned to the published carrot reference genome (Iorizzo et al., 2016) and subsequently filtered for Single Nucleotide Polymorphisms (SNPs), applying default values for all pipeline parameters. After quality filtering, 379 out of 414 Queen Anne’s lace samples and all the cultivated samples (144) remained.

After the initial SNP calling, we merged our dataset with eight re-sequenced samples from Iorizzo et al., (2016), using the merge function of bcftools. Re-sequenced samples included three wild samples from Asia (from Uzbekistan, Turkey, and China), four wild samples from Europe (three from Portugal and one from France), and one sample of *D. syrticus*, a wild carrot species closely related to Queen Anne’s lace, from Tunisia (Arbizu, Ellison, Senalik, Simon, & Spooner, 2016). After merging, the indels were removed and the SNPs were filtered to keep only the biallelic sites, with < 30 % missing data and MAF > 1 % and then individuals with > 40 % missing data were removed. For the population structure analyses (ADMIXTURE, PCA, and DAPC), the SNPs were thinned to 1 SNP per kb. For the introgression analyses (*Dsuite* and *Dinvestigate*), the samples of Queen Anne’s lace from Europe and East were removed, and the sample of *D. syrticus* (accession Ames 29108) was included as an outgroup but the SNPs were not thinned.

### Genetic diversity and population structure

An analysis of molecular variance (AMOVA) was used to partition genetic variation hierarchically between groups (cultivated and wild), between populations within groups, and between individuals within populations, using Arlequin v.3.5.2.2 (Excoffier, Laval, & Schneider, 2005). Three populations formed the cultivated group: CROP_EAST (N = 61), CROP_EUROPE (N = 40), and CROP_US (N = 15); and 29 populations formed the wild group (N = 4 – 20 individuals per population; Table 1). We did not include any samples from the CROP_OTHER category because of their unknown origin (pre-breeding lines) or poor representation (two cultivars from South Africa and one from New Zealand). The statistical significance of genetic differentiation (F_ST_, F_SC_, and F_CT_ for the genetic variance between groups, between populations within groups, and between individuals within populations, respectively), and pairwise F_ST_ were tested nonparametrically after 10,000 permutations. We also used Arlequin to estimate Theta S, as a measure of genetic diversity for each population.

Population structure was analyzed using the thinned dataset with three distinct methods: 1) ADMIXTURE, run from K= 2 - 10, with default parameters (Alexander & Lange, 2011); 2) principal component analysis (PCA), with the R package *SNPRelate* (Zheng et al., 2012); and 3) a discriminant analysis of principal components (DAPC), with the R package *Adegenet* (Jombart, 2008; Jombart, Devillard, & Balloux, 2010). With ADMIXTURE and DAPC, the best K value was determined using cross-validation scores and the Bayesian Information Criterion (BIC), respectively.

### Gene flow between cultivated and wild carrot

To test for gene flow between cultivated and wild carrot, we performed ABBA-BABA tests with *Dsuite* (Malinsky, Matschiner, & Svardal, 2021). We performed these analyses using the 29 wild populations and CROP_US, together with accession Ames 29108 of *D. syrticus* as the outgroup. We first used *Dtrios* with default parameters (Malinsky et al., 2021) to estimate Patterson’s *D* and its associated P-values; together with Z-scores and f4-ratios (an estimate of the fraction of introgression), for all trios of populations (4060 tests). Patterson’s *D* and f4-ratios were calculated from biallelic SNPs across four populations: P1, P2, P3, and the outgroup (O), related by a simple phylogeny (((P1, P2), P3), O) (Malinsky et al., 2021). The allele carried by the outgroup is designated as the ancestral allele (A), while the derived allele is designated as B. Under the null hypothesis, which assumes no gene flow, the ABBA (B shared by P2 and P3) and BABA (B shared by P1 and P3) patterns are expected to occur with equal frequency due to incomplete lineage sorting. A significant increase in ABBA or BABA is consistent with introgression between P3 and either P1 (ABBA < BABA, negative *D* values) or P2 (ABBA > BABA, positive *D* values). As we were interested in the gene flow between crop and wild, we only considered trios with CROP_US as P3 (405 tests). Trios showing Z-scores > 4 were considered statistically significant.

### Additional methods to study the timing and direction of gene flow

ABBA-BABA tests provide evidence of shared alleles between two populations but do not provide any information on either the timing or direction of the gene flow, i.e., positive *D* scores can be produced by crop to wild, wild to crop, or bidirectional gene flow, and by recent and / or historical migration events. As we are interested in the crop to wild gene flow, with populations showing evidence of admixture with the crop, we employed two additional methods to test whether the ABBA-BABA patterns could be explained by the introgression of crop alleles into wild populations. First, we performed demographic modelling with FASTSIMCOAL2 v.2709 (Excoffier et al., 2021) to directly compare scenarios with recent gene flow from crop to wild, wild to crop, bidirectional gene flow, and with no gene flow. FASTSIMCOAL2 uses coalescent simulations to model demographic scenarios from the site frequency spectrum (SFS) (Excoffier, Dupanloup, Huerta-Sánchez, Sousa, & Foll, 2013; Excoffier et al., 2021). For constructing the SFS files, individuals were grouped into three populations, P1 represented wild populations without admixture, P2 included populations with signs of admixture with CROP_US; and P3 comprised the pool of cultivated alleles in the United States (Table S2 for individuals used). To reduce the number of models and the computational effort, we merged all the Massachusetts populations (MA1-MA4) into one (hereafter MA). To avoid biases due to differences in sample sizes, we included the same number of individuals for P1 and P2 (Table S2). Also, as the computation of SFS assumes independence between loci and is sensitive to missing data, the datasets were filtered to retain individuals with little missing data (< 20 %), and SNPs with MAF > 0.05 and low linkage disequilibrium (1 SNP per kb). Multidimensional minor allele site frequency spectrum files were produced using ARLEQUIN and were used as the input file in FASTSIMCOAL2.

Four models varying in migration events between P2 and P3 were compared (M1: no migration; M2: migration from crop to wild; M3: migration from wild to crop; and M4: bidirectional migration). In all models, we fixed the timing of divergence between crop and wild (i.e., domestication; *T_DIV1_*) to 1100 generations ago, whereas the other parameters were estimated by the model: migration parameters (*m_CW_* and *m_WC_* for crop to wild and wild to crop, respectively), current population sizes of P1, P2, and CROP (*N_P1_, N_P2_*, and *N_CROP_*, respectively), timing of divergence between P1 and P2 (*T_DIV2_*), and the bottleneck intensity at *T_DIV1_*. All migration events were assumed to have occurred after the divergence of P1 and P2. Each model (M1-M4) was run 50 times, each with 100,000 simulations and 40 optimization cycles. For each run, we obtained the Akaike information Criterion (AIC = −2 * lnL + 2 * K), where lnL is the likelihood provided by the model, and K is the number parameters in the model. To identify the best model, we made pairwise comparisons with Mann-Whitney U tests. After that, to parameterize the best model, point estimates of parameters were taken from the best run, while minimum and maximum values for parameters were obtained from the best five runs. An example of files used for simulations can be found in Supplementary file 1.

Finally, using the same samples but incorporating *D. syrticus* as the outgroup, we estimated the *D* frequency spectrum (*D_FS_*) (Martin & Amos, 2021). The *D_FS_* is an extension of the ABBA-BABA tests which partitions the introgression signs into allele frequency bins, thus providing information on the timing and direction of introgression events (Martin & Amos, 2021). For recent introgressions, we expect positive *D* scores to be distributed mostly in the lower allele frequency bins (top-left in the plots), while for older introgressions positive *D* scores should occur in the intermediate and high allele frequency bins (top-right in the plots).

### Identification of introgressed regions

For populations with evidence of recent crop to wild introgression (CA1, CA4, MA1 - MA4), we used *Dinvestigate* (Malinsky et al., 2021) to localize introgressed regions along the genome and to test whether introgression was confined to specific loci or distributed throughout the genome. For each population, we calculated the statistic *distance fraction* in non-overlapping windows of 20 informative SNPs, i.e., SNPs with either the ABBA, BABA, or BBAA pattern (Malinsky et al., 2021; Pfeifer & Kapan, 2019). This statistic uses a genetic distance-based approach to estimate introgression, which is more robust than Patterson’s *D* for smaller genomic regions (Malinsky et al., 2021; Pfeifer & Kapan, 2019). Briefly, with the phylogeny (((P1, P2), P3), O), the statistic *distance fraction* is the difference between the genetic distances between P1 and P3 (*D_13_*) and between P2 and P3 (*D_23_*). For each window, the *distance fraction* varies from 1 to −1; positive values indicate an admixture between P3 and P2 (*D_23_* < *D_13_*) and negative values an admixture between P3 and P1 (*D_23_* > *D_13_*). We used all the individuals from Iowa as P1, all individuals for either CA1, CA4 or MA1-MA4 as P2, all cultivars from the United States (CROP_US) as P3, and *D. syrticus* as the outgroup (O). Individuals from Iowa were used as P1 because the IA populations showed no evidence of admixture with the crop, and low genetic differentiation with the P2 populations. Without introgression, *D_23_* and *D_13_* are similar and the distance fraction values are close to 0. The top 5% windows (higher *distance fraction* values) were considered significant. The SNPs within the introgressed regions identified were compared between populations from the same region (i.e., California or Massachusetts), using Venn diagrams.

## RESULTS

### Sample and SNP information

After filtering, 523 samples (144 cultivated + 379 wild; Tables S1 and S3) and 34,871 SNPs (full dataset) were kept. After thinning to 1 SNP per kb, 13,073 SNPs remained (thinned dataset). Genetic diversity estimates and population structure analyses (ADMIXTURE, PCA, and DAPC) were performed with the thinned dataset, whereas the full dataset was used for the ABBA-BABA tests.

### Genetic diversity and population structure

With AMOVA, we observed significant differences between groups (cultivated vs. wild; F_ST_ = 0.209; P < 0.001), between populations within groups (F_SC_ = 0.095; P < 0.001), and within populations (F_CT_ = 0.126; P < 0.001). Genetic diversity (Theta S) for cultivated groups varied from 1401 ± 428 (CROP_US) to 1452 ± 363 (CROP_EUROPE), and from 854 ± 280 (CO) to 2196 ± 737 (IA3) between wild populations (Table S4). All wild populations but one (CO) showed higher genetic diversity than the cultivars (Table S4). Pairwise F_ST_ indicated an overall low population structure among the wild populations, and similar genetic differentiation with cultivars (Table S4).

ADMIXTURE clearly differentiated cultivated and wild samples (Fig. 1). At K = 2, 122 / 144 and 135 / 144 cultivated samples have a K1 ancestry > 0.99 and > 0.90, respectively (Fig. 1), suggesting no recent introgression of wild genomic regions into the cultivated germplasm. In contrast, many wild samples showed evidence of admixture with crop; 93 / 376 wild samples from the United States showed a K1 ancestry > 0.1, ranging from 0.11 to 0.33 (Fig. 1A, 1B). The admixed individuals (K1 > 0.1) mostly belonged to three geographic areas: California (CA) (45 / 69), Colorado (CO) (11 / 11), and Massachusetts (MA) (31 / 39). At K = 3, the cultivated samples remained assigned to a single genetic cluster and the wild samples were split into two genetic clusters: one of the clusters included samples from Wisconsin (WI), Iowa (IA), and Minnesota (MN), and the other included samples from CO and CA4 (Fig. 1C). Samples from CA (all except CA4), South Dakota (SD), and MA populations seemed to be admixed between these two genetic clusters (Fig. 1C). At K = 4 samples from CO became a group on their own (Fig. 1C), and at K = 5 cultivated samples were split into two genetic clusters, mostly differentiating East cultivars from European and US cultivars (Fig. 1C). At higher K values, geographic regions and then individual populations became groups on their own (Fig. S1).

**Figure 1.**
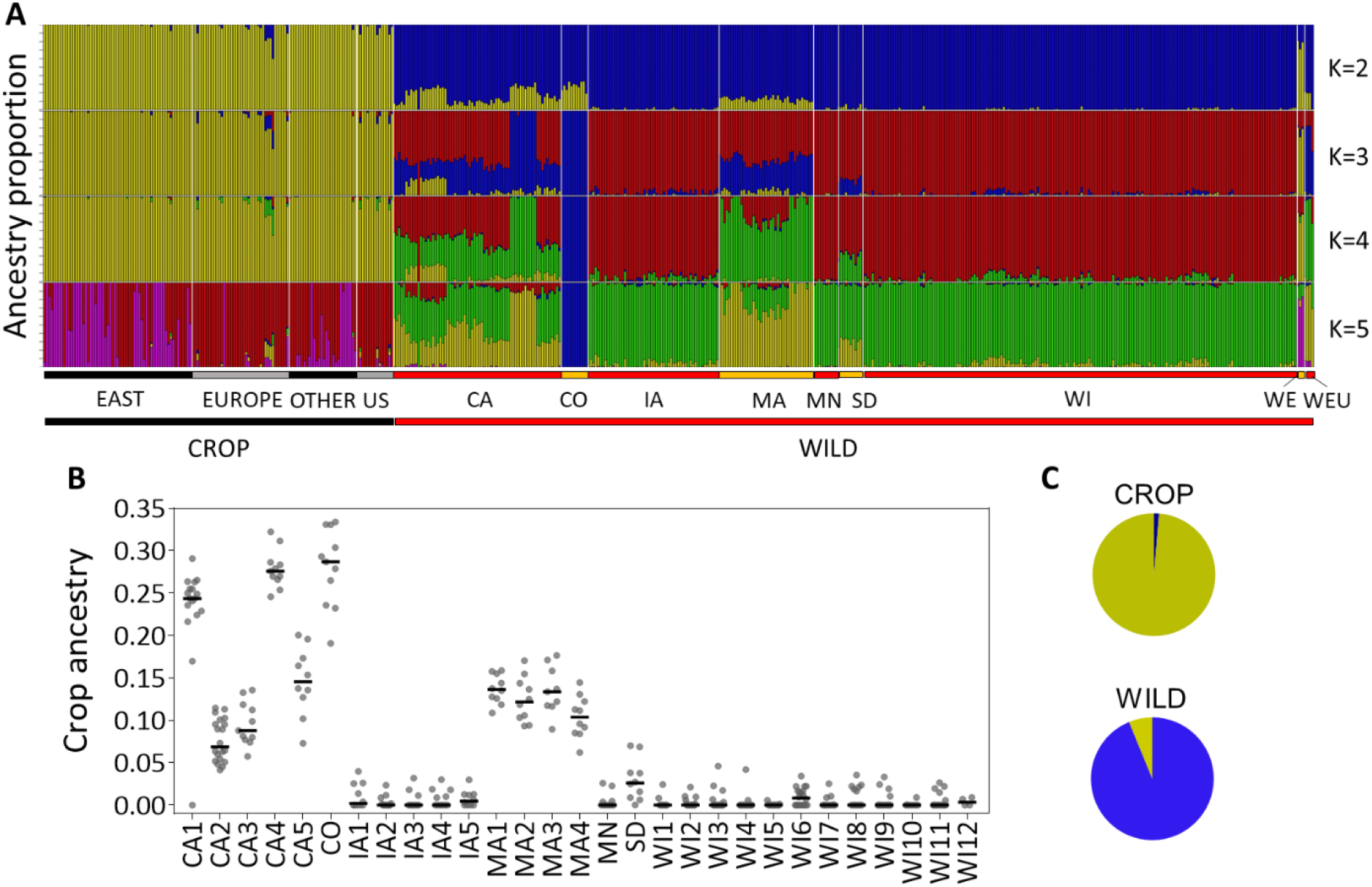
Admixture proportions and Crop ancestry. A) Ancestry proportions at K = 2 - 5. Cultivars are grouped by region, and wild individuals are grouped by state. B. Crop ancestry (at K = 2) of individuals grouped by populations. C. Pie charts showing mean ancestry for the crop and wild groups. Information for samples can be found in Tables S1 and S3.

Principal component analysis recapitulated the results seen in ADMIXTURE. The first axis (PC1) separated the cultivated and wild samples, and the second axis (PC2) split the cultivated samples into two genetic clusters (Fig. 2A), mostly corresponding to those identified with ADMIXTURE. The wild samples from Europe (three from Portugal and one from France) grouped with the wild samples from the United States, while the three wild samples from Asia (Wild_East, from Uzbekistan, Turkey, and China) grouped with the East cultivars (Fig. 2A). Principal component 3 and PC4 separated samples from CO and CA4 from the rest, respectively (Fig. 2B), and the wild accession from France (PI 478861) grouped with CA4 (Fig. 2B). Clustering analysis with DAPC suggested three groups (Figs. 2C and 2D), largely consistent with PCA (Fig. 2A). Pairwise F_ST_ indicated that the three DAPC groups were well differentiated (Fig. 2E). When the wild samples from the United States were analyzed separately, similar results were observed in terms of population structure, with CO and CA4 well differentiated from each other, and with the rest of the populations (showing an overall low population structure) located in between CO and CA4 (Fig. S2).

**Figure 2.**
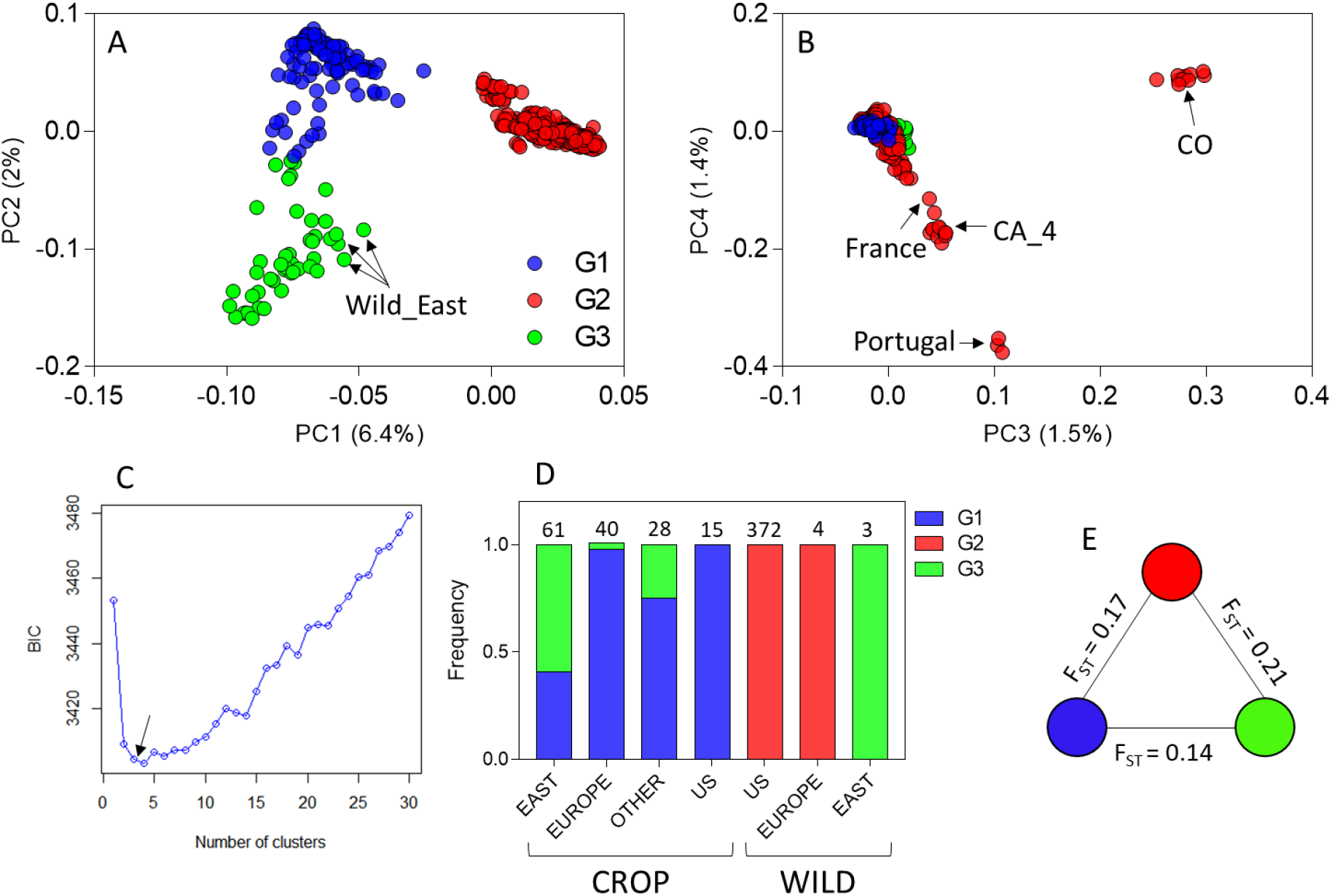
Principal component analysis (A and B) and discriminant analysis of principal components (DAPC) (C and D) of wild and cultivated samples of carrot. In A and B, individuals are colored by their membership to DAPC groups. C. The relationship between Bayesian Information Criterion (BIC) and the number of clusters supports the presence of three groups. D. Bar plot showing the frequency of individuals assigned to each DAPC group, the number of individuals is indicated above the bar. E. Pairwise F_ST_ between DAPC groups. Information for samples and membership of DAPC groups can be found in Tables S1 and S3.

### Gene flow between cultivated and wild carrot

Using ABBA-BABA tests, we found significantly positive Patterson’s *D* scores (Z-scores > 4) in 140 / 405 tests involving CROP_US as P3 (Table S5). Seven (from three states) out of 29 populations showed significant admixture with CROP_US (Tables 3 and S5). The fraction of the introgressed genome (f4-ratio) varied between populations from 0.152 ± 0.003 to 0.243 ± 0.002 (Table 2). The population used as P1 did not affect the estimation of the fraction introgressed, as denoted by the small standard errors. These results were both qualitatively and quantitatively consistent with the admixture proportions estimated with ADMIXTURE (Table 2).

**Table 2.**
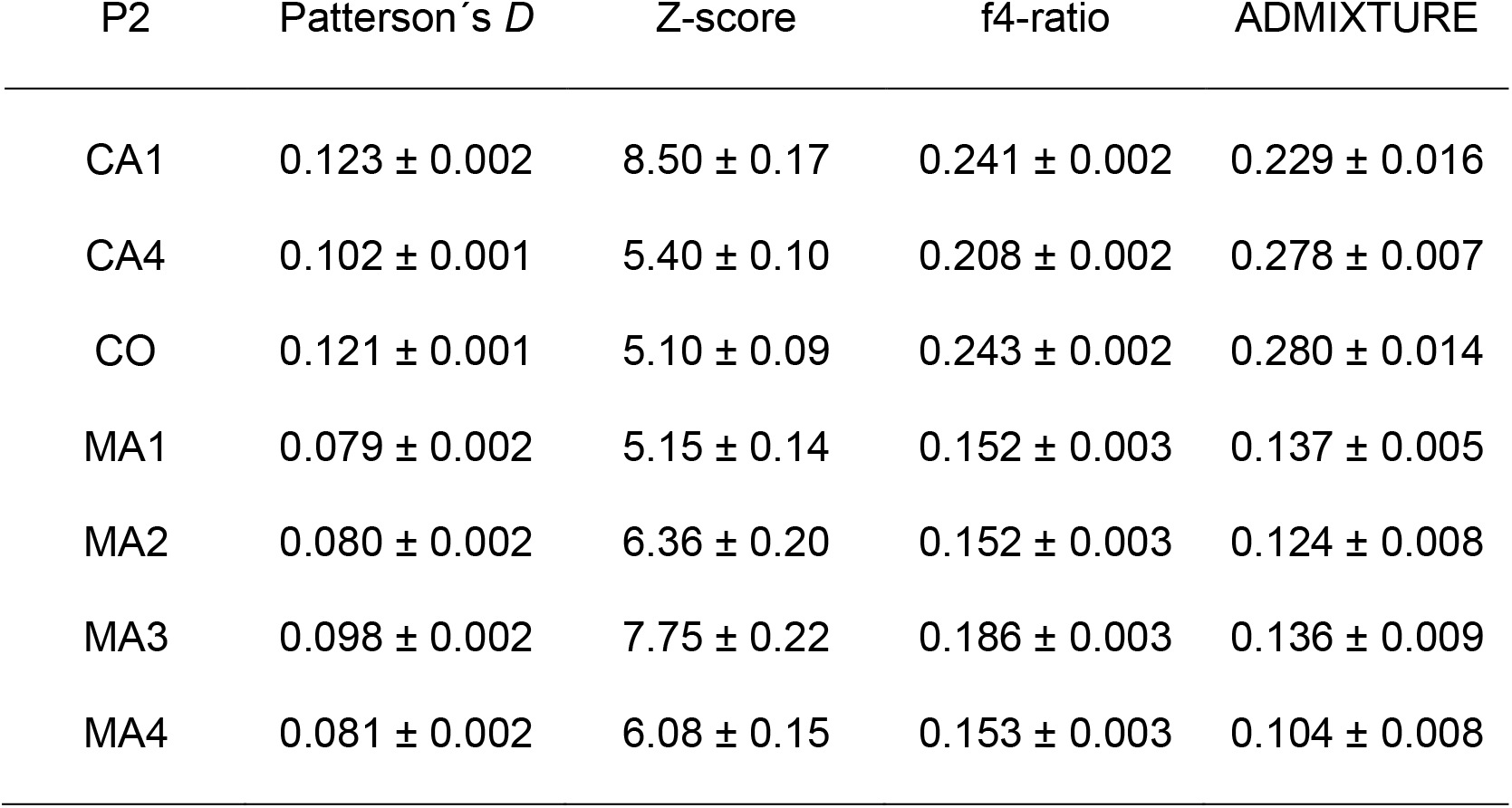
Patterson’s *D* statistic, Z-scores, and f4-ratios for the seven populations (P2) showing significant evidence of crop introgression. To estimate means ± standard errors we used values from the 17 trees involving populations from Iowa (5) and Wisconsin (12) as P1. In all tests, CROP_US was used as donor and one sample of *Daucus syrticus* as outgroup. Cultivated ancestry proportion from admixture at K=2 is included for comparison.

| P2 | Patterson's $D$ | $Z$-score | $f_4$-ratio | ADMIXTURE |
| --- | --- | --- | --- | --- |
| CA1 | $0.123 \pm 0.002$ | $8.50 \pm 0.17$ | $0.241 \pm 0.002$ | $0.229 \pm 0.016$ |
| CA4 | $0.102 \pm 0.001$ | $5.40 \pm 0.10$ | $0.208 \pm 0.002$ | $0.278 \pm 0.007$ |
| CO | $0.121 \pm 0.001$ | $5.10 \pm 0.09$ | $0.243 \pm 0.002$ | $0.280 \pm 0.014$ |
| MA1 | $0.079 \pm 0.002$ | $5.15 \pm 0.14$ | $0.152 \pm 0.003$ | $0.137 \pm 0.005$ |
| MA2 | $0.080 \pm 0.002$ | $6.36 \pm 0.20$ | $0.152 \pm 0.003$ | $0.124 \pm 0.008$ |
| MA3 | $0.098 \pm 0.002$ | $7.75 \pm 0.22$ | $0.186 \pm 0.003$ | $0.136 \pm 0.009$ |
| MA4 | $0.081 \pm 0.002$ | $6.08 \pm 0.15$ | $0.153 \pm 0.003$ | $0.104 \pm 0.008$ |

### Demographic modelling supports crop to wild gene flow

The most likely demographic model for three (MA, CA1, and CA4) out of four P2 populations was model 2 (crop to wild gene flow), followed by model 1 (no gene flow) (Fig. 3; Table S6). For CO, the most likely model was model 3 (wild to crop gene flow), followed by model 1 (Fig. 3; Table S6). Similarly, estimates of *DSF* supported crop to wild gene flow for three out of four comparisons (CA1, CA4, and MA; Fig. 4A, 4B, and 4D). For these populations, introgressed alleles were distributed across frequency bins, being consistent with the ongoing gene flow patterns. In addition, in CA_4, there was a large proportion of shared variants at high frequency (Fig. 4B) suggesting some historical gene flow. For CO, both the negative *DSF* at low frequencies and the large proportion of shared variants at high frequency (Fig. 4C) are consistent with a severe bottleneck and historical gene flow.

**Fig. 3.**
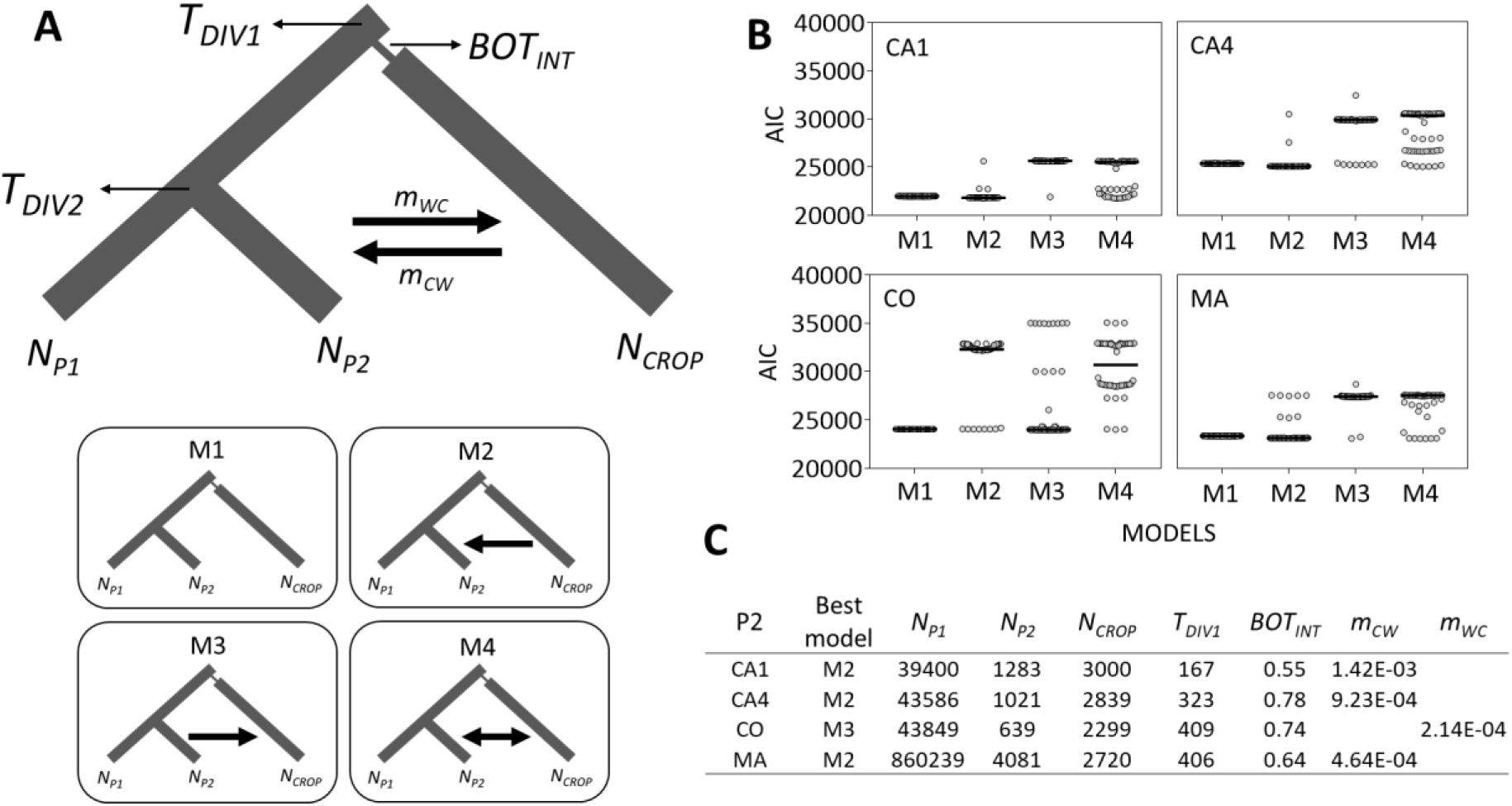
A. Demographic scenarios modelled in FASTSIMCOAL2. For each wild population where introgression was previously detected, four models differing in migration events were evaluated. M1: no gene flow; M2: gene flow from CROP to P2 (*m_CW_*); M3: gene flow from P2 to CROP (*m_WC_*); and M4: bidirectional gene flow (including both *m_CW_* and *m_WC_*). Time of divergence*T_DIV1_* was fixed at 1100 generations ago. The other parameters were estimated for each model: *T_DIV2_*: time of divergence between P1 and P2; *BOT_INT_*: bottleneck intensity, measured as the inverse of the bottleneck size during the bottleneck (1 generation); *N_P1_*: population size of P1; *N_P2_*: population size of P2; *N_CROP_*: population size of CROP. B. Model support based on AIC values for 50 runs, pairwise medians, denoted as horizontal lines, were compared with Mann-Whitney U tests and all comparisons showed statistically significant differences (P < 0.0001). C. Model parameters for the best of run of each model.

**Figure 4.**
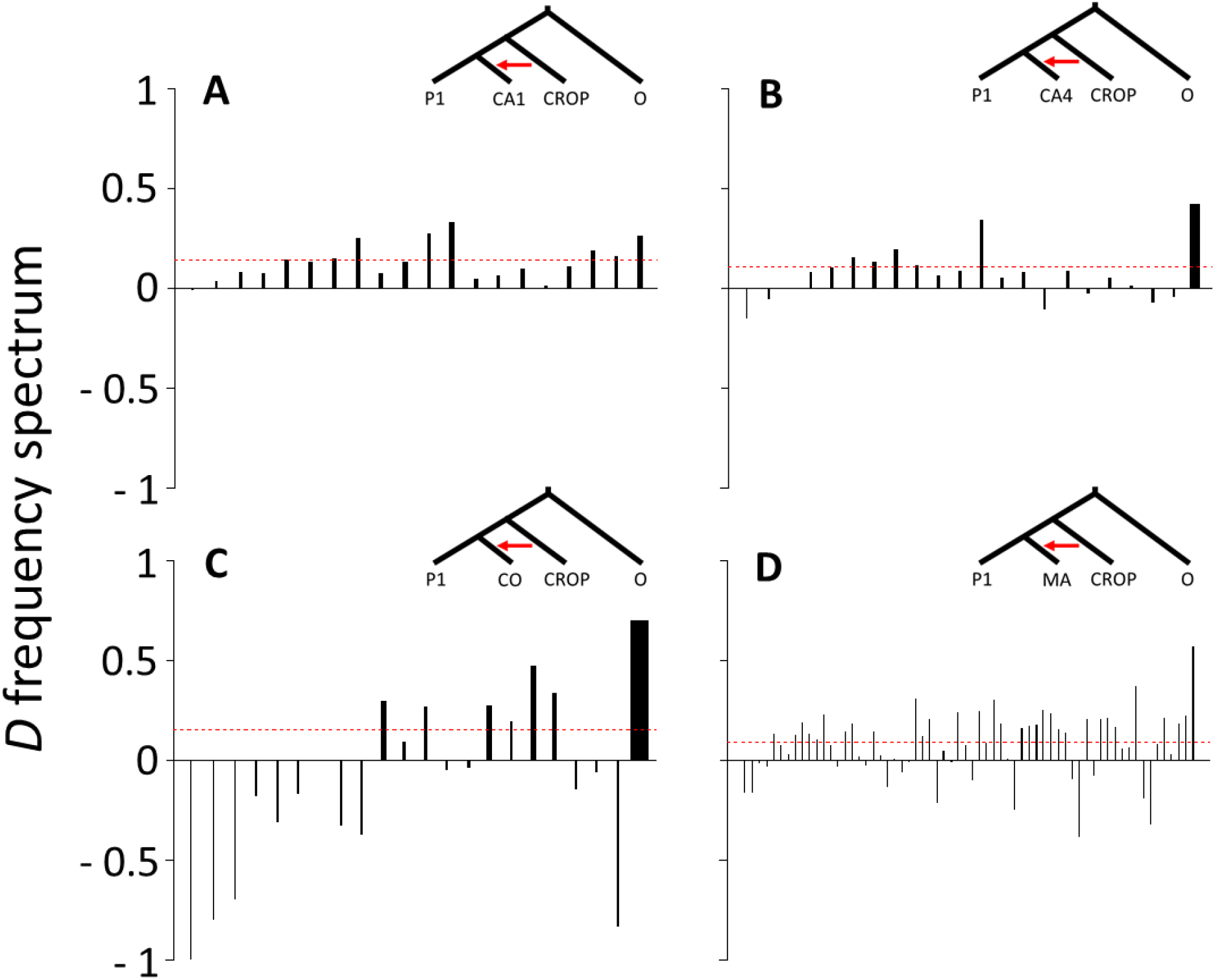
Estimates of the *D* frequency spectrum (*D_FS_*) for populations showing significant results for the genome-wide ABBA-BABA test. A. CA1; B. CA4; C. CO, and D. MA. Phylogenetic trees indicate the most probable demographic model. P1: individuals from Iowa, Wisconsin, and Minnesota. Samples used in each comparison are presented in Table S2.

### Genes introgressed from cultivated carrot are distributed throughout the genome of wild carrot

Introgressed windows were defined as those from the top 5% *distance fraction* values. In all populations, introgressed windows were distributed across all nine chromosomes, and no clear peaks were identified (Fig. 5A). Of the 3369 SNPs located within introgressed windows in six populations, no SNPs were found in all the populations, and there was very little overlap of introgressed SNPs between the MA (5 / 2252) and CA (82 / 1506) populations (Fig. 5B).

**Figure 5.**
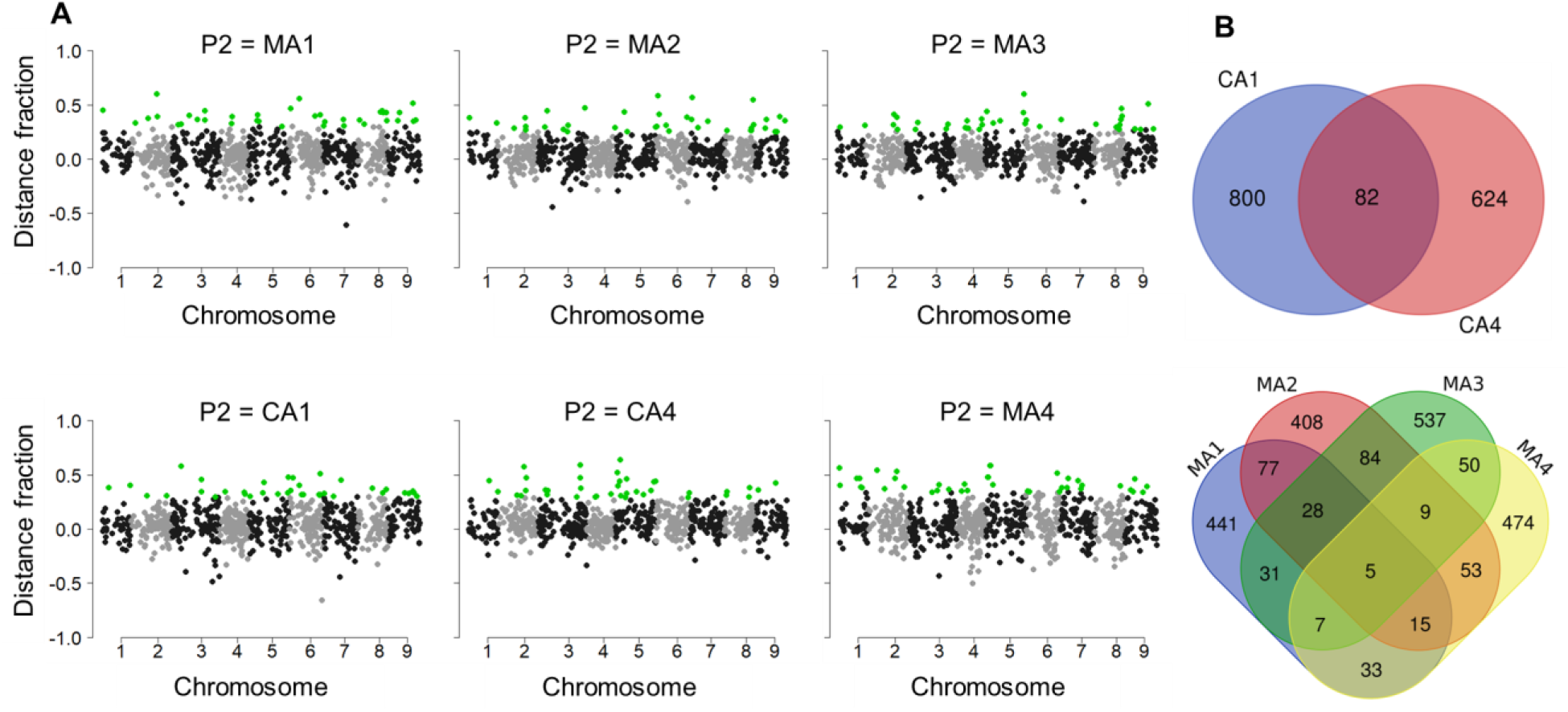
A. Manhattan plots of distance fraction values for populations with the strongest signs of crop introgression. Each dot represents a window of 20 SNPs. Positive and negative values indicate shared ancestry between P3 (CROP_US) and P2 (CA1, CA4, or MA1-MA4) and between P3 and P1 (all individuals from IA), respectively. Top 5% outliers highlighted in green. In all cases, one sample of *Daucus syrticus* was used as outgroup. B. Venn diagrams for SNPs within introgressed windows.

## Discussion

We investigated whether wild carrot populations in the United States carry crop alleles as a consequence of crop to wild introgression. Results of the population structure analyses are consistent with what we knew about the evolutionary history of cultivated and wild carrot in the United States (Iorizzo et al., 2016, 2013). Cultivars and wild populations form two well differentiated groups, and both genetic groups are, respectively, genetically similar to their European counterparts. In Europe, cultivated and wild carrots hybridize naturally (Magnussen & Hauser, 2007), and, therefore, hybridization could have occurred prior to the introduction of wild carrot to the United States (Iorizzo et al., 2013). We detected admixture in several, but not all, wild carrot populations. This pattern suggests either post introduction admixture, or multiple introductions, with introduction of wild carrot in some regions, and of crop-wild hybrids (admixed) in others. However, we identified low divergence between admixed and non-admixed populations, which supports post introduction admixture. Hybridization between crops and their wild relatives from outside their native range has previously been reported for other crop-wild complexes, such as maize - teosinte in Europe (Le Corre et al., 2020), bread wheat - jointed goatgrass in the United States (Martins, Sun, & Mallory-Smith, 2015), and cultivated - prairie sunflower in Argentina (Mondon et al., 2018).

Though ADMIXTURE and ABBA-BABA tests are robust methods to detect admixture between populations or related species, such methods provide little or no information on the timing and direction of introgression (Hibbins & Hahn, 2022; Martin & Amos, 2021; Payseur & Rieseberg, 2016). Therefore, we used additional methods to study the direction and timing of gene flow. Two approaches (demographic modeling in FASTSIMCOAL2, and *D_FS_*) support a direction of introgression from cultivars to wild populations in California and Nantucket, but hybridization history is less clear in Colorado.

California is the state with the highest carrot production in the United States; mainly as root crops but there is also some seed production in areas where wild carrots are absent or very rare. In Nantucket, carrots are grown as root crops. In fields for root production, the plants are harvested before flowering, and therefore, one would expect little to no gene flow between the crop and wild populations. However, we found significant amounts of crop ancestry in most wild individuals from these areas. Similarly, a previous study on wild carrot populations in Nantucket using SSR markers found that the populations closer (< 1 Km) to crop fields were less differentiated with the crop than populations further away (> 2.5 Km), suggesting crop to wild introgression in the vicinity of crop fields (Mandel et al., 2016). Here, we used populations collected at least 2.5 km away from crop fields and found consistent signs of crop introgression, demonstrating that crop alleles have spread across the island. In addition, in California, the wild carrot populations sampled were not in close proximity to the main carrot production areas (Calflora, 2022), and no carrot fields were visually identified in the vicinity at collection (A. Flick and P. Simon, pers. com.). Therefore, if crop and wild were allopatric, it is unclear where hybridization could have taken place. A possible explanation is that hybridization took place near carrot production areas, and then cultivar alleles spread via gene flow. This scenario is supported by high rates of outcrossing and long-distance pollen dispersal by insects in carrots (Rong et al., 2010).

In Colorado, demographic modeling did not support crop to wild introgression. This population has experienced a severe bottleneck, as shown by the very low genetic diversity (~ 40 % relative to other wild populations: Table S3), and the distribution of *D_FS_* across allele frequencies (Fig. 4). The *D_FS_* analysis shows that most shared variants are fixed in CO, which is more consistent with the historical than recent gene flow (Martin & Amos, 2021). Therefore, processes other than introgression, such as incomplete lineage sorting and genetic drift, probably resulted in an increase of shared variants between CO and cultivars (Hibbins & Hahn, 2022; Martin & Amos, 2021). This case highlights the importance of characterizing hybridization events, especially in studies of crop-wild gene flow, where the timing and direction of gene flow have direct implications on risk assessment.

### Are crop alleles neutral in admixed wild carrot populations?

Crop alleles introduced in recipient wild populations can be adaptive, deleterious, or neutral. Each case leaves a distinct signature on the genome (Hufford et al., 2013; Moran et al., 2021; Runemark, Vallejo-Marin, & Meier, 2019) which can be interpreted with the use of current population genomic methods (Hibbins & Hahn, 2022; Martin, Davey, & Jiggins, 2015; Payseur & Rieseberg, 2016). For example, due to selection and recombination, adaptive introgressed variants increase in frequency while surrounding variants are removed by negative selection (deleterious variants) and/or by genetic drift (neutral variants), creating peaks identifying adaptive introgression across the genome (Burgarella et al., 2019; Rendón-Anaya et al., 2021). In this study, we observed no such peaks, but individuals from admixed populations carried multiple, small genomic regions from cultivars, with very little overlap between populations from the same geographic origin (Fig. 5), suggesting that most crop alleles were neutral. Neutrality of crop alleles in carrot is consistent with phenotypic studies showing that crop-wild hybrids can perform as well as wild carrots (non-hybrids) (Hauser & Shim, 2007). Our results also suggest that the neutrality of crop alleles is environment-dependent, i.e., alleles that are neutral in California and Nantucket may be deleterious in other areas such as Wisconsin, where hybridization probably occurs (Palmieri et al., 2019) but introgression does not proceed.

In both regions where crop introgression was detected, California and Nantucket, we did not identify any pure wild carrot individuals (except one in CA1; Fig. 1B). As wild carrot has been present in these regions since at least the early 1900s (Owens 1888; Calflora 2022), this pattern suggests that cultivar alleles have spread across wild populations under neutrality, replacing pure wild genomes. Theory predicts that when the fitness of hybrids is not strongly reduced (as has been shown in carrot), the frequency of hybrid individuals in a population can increase over time without the genetic diversity being reduced (Hedge et al., 2006; Runemark et al., 2019; Todesco et al., 2016). As predicted, we observed no reduction of genetic diversity in the admixed populations, nor evidence of bottlenecks. Our finding that most introgressed crop alleles seem to be neutral in these populations supports this scenario of introgression under neutrality. Genotyping herbarium specimens from these regions could shed light on the temporal dynamics of crop allele introgression and persistence.

### Do environmental variables play a role in crop introgression?

Despite clear evidence of introgression and persistence of crop alleles in California and Nantucket, we did not find any evidence of crop introgression in 19 populations (out of 29) collected in four states (Iowa, Minnesota, South Dakota, and Wisconsin). While allopatry with the crop may explain the absence of introgression in some areas, we did not find evidence of introgression in well-known sympatric populations from Madison, Wisconsin (WI1 – WI4). These populations were collected less than 500 m away from a site that has been used to breed carrot cultivars for over 40 years, and we therefore suspect post-hybridization barriers could be preventing crop allele introgression (Campbell, Snow, & Sweeney, 2009; Presotto et al., 2019). In this study, the admixed populations tended to occur in areas with milder winters than the non-admixed populations (Table 1; Fig. S4), suggesting that harsher winters may prevent crop introgression. In wild carrot, overwinter survival is a critical stage in its life history, strongly affecting population growth (Hauser & Shim, 2007; Van Etten & Brunet, 2017). As carrot crops are not bred to withstand harsh winters, hybridization with crop could negatively affect the overwinter survival of hybrid individuals. Hauser (2002) reported that at – 4 °C crop-wild hybrids survived as well as the wild individuals, but at – 8 °C the survival of hybrids was lower than for wild carrot plants. Therefore, in regions (and seasons) with milder winters, such as California, crop-wild hybrids may overwinter and survive to reproduce (Hauser & Shim, 2007), whereas in regions with harsher winters, such as Wisconsin, crop-wild hybrids may not survive the winter due to their lower frost tolerance, preventing introgression. Selection experiments comparing crop, wild, crop-wild hybrids and backcrosses, in different regions (e.g., in Nantucket, California, Wisconsin and Iowa) would further our understanding of the impact of environmental conditions on the introgression of crop alleles into wild carrot populations.

### Implications

In many hybridization studies, some regions of the genome are found to be intolerant to introgression while others are more prone (Hanemaaijer et al., 2018; Hufford et al., 2013; Moran et al., 2021). In the current study, we did not observe any such pattern and cultivar alleles were distributed throughout the genome. If certain regions of the genome were intolerant to introgression, the introduction of new genes or edited genes into these “introgression intolerant” chromosome regions could be a strategy to limit the introgression of GM genes into wild populations, However, this strategy has never been proven in any crop species and would not be viable in carrot.

Since the fate of hybrids can be environmentally dependent (Moran et al., 2021), one would expect the changes in environmental conditions resulting from climate change to impact introgression, although the evidence supporting such assertation remains weak (Todesco et al., 2016). Our results suggest that warmer environments are more suitable for crop introgression, thus, milder conditions resulting from climate change could increase the potential survival of hybrid individuals and the incidence of introgression.

Based on the results of this study, we anticipate that the release of genetically engineered carrot cultivars would lead to the introduction and spread of cultivar alleles in wild carrot populations, at least in regions with milder winters. Similar questions should be addressed in other crops with wild or feral relatives to further quantify the impact of the new genome-editing technology on the introgression of cultivar genes into wild or feral populations.

## Conclusions

Using multiple approaches to detect and characterize introgressions, we identified two regions of the United States where cultivar alleles had introgressed into wild carrot populations: California and the Nantucket Island. In these areas, no pure wild genotypes were found, indicating that crop alleles may quickly spread in wild populations. Admixed populations were found in environments with milder winters than non-admixed populations, suggesting influence of the environment on crop introgression. We did not identify any large genomic region intolerant to introgression, and we found no support for adaptive (nor maladaptive) introgression, instead, most crop alleles seem to be neutral for admixed populations.

## Supplementary material

Table S1. Information of cultivated accessions used in this study.

Table S2. Samples used for demographic modeling in FASTSIMCOAL2.

Table S3. Information of wild samples used in this study.

Table S4. Genetic diversity and population structure of wild and cultivated carrots.

Table S5. Gene flow models (ABBA-BABA tests) for all trios of populations.

Table S6. Parameter estimates and Akaike Information Criterion (AIC) for all runs and models implemented in FASTSIMCOAL2.

Supplementary files 1. Example of files used for simulations in FASTSIMCOAL2.

Fig. S1. Admixture plots for K = 6 - 10.

Fig. S2. Principal component analysis of wild samples from the United States.

Fig. S3. Variation in mean annual temperature (°C) (BIO1), minimum temperature of the coldest month (°C) (BIO6), and annual precipitation (mm) (BIO12) between admixed and non-admixed populations.

## Acknowledgements

We thank Andrew Flick and other members of the Brunet Lab for collecting samples; Jennifer Mandel for providing the Nantucket samples; and Philipp Simon for collecting some California samples and providing samples of pre-breeding lines from the Wisconsin Breeding Program.

We thank the National Research Council of Argentina (CONICET) for a fellowship to FH, and the United States Department of Agriculture, Agricultural Research Service for providing funding to JB for this project. LP was on an appointment with the Agricultural Research Service (ARS) Research Participation Program administered by the Oak Ridge Institute for Science and Education (ORISE) through an interagency agreement between the U.S. Department of Energy (DOE) and the U.S. Department of Agriculture (USDA).

## Author Contributions

Johanne Brunet conceived and oversaw the project. Luciano Palmieri extracted DNA and processed the sequence data. JB and LP participated in sample collection. Fernando Hernández conducted the data analyses and wrote the manuscript with contributions from all authors.

